# ESD1 Affects Seed Setting Rate in Rice by Controlling Embryo Sac Development

**DOI:** 10.1101/2021.02.01.429182

**Authors:** Tiankang Wang, Yixing Li, Shufeng Song, Mudan Qiu, Licheng Zhang, Chengxia Li, Hao Dong, Lei Li, Jianlong Wang, Li Li

**Affiliations:** State Key Laboratory of Hybrid Rice, Hunan Hybrid Rice Research Center, Changsha 410125, China; College of Agronomy, Hunan Agricultural University, Changsha 410128, China

**Keywords:** rice, ESD1, seed setting rate, embryo sacs

## Abstract

Seed setting rate is one of the critical factors that determine rice yield. Grain formation is a complex biological process, whose molecular mechanism is yet to be improved. Here we investigated the function of an OVATE family protein, Embryo Sac Development 1 (ESD1), in the regulation of seed setting rate in rice (*Oryza sativa*) by examining its loss-of-function mutants generated via CRISPR/Cas9 technology. *ESD1* was predominantly expressed at stage 6 of panicle development, especially in the ovules. *esd1* mutants displayed reduced seed setting rates with normal stamen development and pollen tube growth but abnormal pistil group. Investigation of embryo sacs revealed that during the mitosis of functional megaspores, some egg cells degraded during differentiation in *esd1* mutants, thereby hindering subsequent fertilization process and reducing seed setting rate. In addition, the transcriptional level of *OsAPC6,* a reported embryo sac developing gene, was found to be significantly reduced in *esd1* mutants. These results support that ESD1 is an important modulator of embryo sac development and seed setting rate in rice. Together, this finding demonstrates that ESD1 positively regulates the seed setting rate by controlling embryo sac development in rice, and has implications for the improvement of rice yield.

## Introduction

Rice is one of the most important food crops in the world. At present, the controlling mechanism of rice yield has made great progress, particularly in grain weight and effective panicles, but little is known about the seed setting rate. Seed setting rate is affected by several factors, including biomass-spikelet ratio, spikelet-to-leaf ratio, specific leaf weight, spikelet-sheath quantity, and spikelet differentiation (Zhou et al., 2011; Toh et al., 2012; Li et al., 2013; Xu et al., 2017; Liu et al., 2018; He et al., 2019; Liu et al., 2019; Xu et al., 2019; Xiang et al., 2019). Moreover, seed setting rate is also easily affected by the external environment, such as drought, high temperature, low temperature, or saline-alkali stress (Xu et al., 2014; Fu et al., 2014; Wang et al., 2016b; Ma et al., 2018; Cheabu et al., 2018; Si et al., 2018). Further study of seed setting rate has important significance both for its mechanism and for the improvement of rice yield.

As an essential part of spikelet, embryo sac development can directly affect seed setting rate. The development of rice embryo sacs can be divided into two stages, the first stage involves the formation of functional megaspores, in which the megasporocyte undergoes meiotic division to yield four haploid cells, three of which disappear, while one becomes the functional megaspore, the second stage is the formation of embryo sac, in which the functional megaspores undergo mitosis three times and eventually form a mature embryo sac, there have been some reports about genes related to embryo sac development (Zee 1997; Li et al., 2007; Chang et al., 2009; Wang et al., 2012; Hou et al., 2019; Khanday et al., 2019; Wang et al., 2019; Xu et al., 2020). OsRAD51C, which belongs to the RecA/RAD51 family, is associated with the normal meiotic division of megasporocyte and microsporocyte at an early stage (Kou et al., 2012; Tang et al., 2014). OsMSH4 and OsMSH5, the members of the ZMM family, can form a heterodimer, which plays a critical role in the regulation of homologous chromosome exchange in the meiotic division (Luo et al., 2013; Zhang et al., 2014). Especially, OsMSH4 influences the exchange of homologous chromosomes during meiotic division of male and female (Wang et al., 2016a). The *OsAPC6* gene encodes an APC6 protein, affecting the development of female gametes but not the development of male gametes. OsAPC6 is crucial for the smooth transition from mitosis metaphase to mitosis anaphase. The T-DNA inserted *osapc6* mutants are fertile in males and partially sterile in females. The plants exhibit dwarfed, and their seed setting rates are reduced by 45% (Kumar et al., 2010; Awasthi et al., 2012).

The OVATE family proteins (OFP) are plant-specific proteins with a conserved OVATE domain (Liu et al., 2014). The discovery of OFP protein family originated from the gene *OVATE* which regulates the shape of tomato fruit. The C-terminal of OVATE protein contains a highly conserved OVATE domain (Liu et al., 2002). The genes of OFP protein family are widely presented in various plant species such as Arabidopsis *(Arabidopsis thaliana),* rice *(Oryza sativa),* tomato *(Solanum lycopersicum),* and pepper *(Capsicum annuum).* OFP family proteins play a key role in regulating the growth and development of plants, including the construction of fruit shape, fruit maturity and quality formation, DNA repair, vascular bundle development, secondary cell wall formation, and embryo sac development (Hackbusch et al., 2005; Wang et al., 2008; Pagnussat et al., 2007; Wang et al., 2010; Li et al., 2011a; Toh et al., 2012; Schmitz et al., 2015; Yang et al., 2016; Ma et al., 2017; Xiao et al., 2017).

In our previous study, microarray data showed that ESD1 protein belongs to the OFP family was highly and abundantly expressed mainly in spikes. In this study, we find that ESD1 positively regulates the rice seed setting rate by controlling embryo sac development. Its loss-of-function *esd1* mutants were successfully generated via CRISPR/Cas9 technology. Results showed that *esd1* mutants exhibited abnormal development of embryo sac, thereby ultimately decreasing the seed setting rate. Meanwhile, the transcriptional level of *OsAPC6*, an embryo sac development related gene, was found to be significantly reduced in *esd1* mutants. *ESD1* had the highest level of expression at stage 6 of young panicle development, especially in the ovules, and ESD1-GFP fusion proteins predominantly localized in cytoplasm. Our results support ESD1 as a key modulator of rice seed setting rate. This finding has significance for the improvement of rice yield.

## Results

### ESD1 belongs to the OFP family

*ESD1* contains only one exon without intron, and its predicted protein ESD1 has 253 amino acids (aa). The SMART database (Simple Modular Architecture Research Tool, http://smart.embl-heidelberg.de/) showed that ESD1 contains an OVATE domain located at 164-227 aa, and belongs to the OFP family (Hackbusch et al., 2005; Liu et al., 2014) (Fig. 1A). The OFP family has 33 members in rice (from OsOFP1 to OsOFP33), and ESD1 is numbered OsOFP31 (Liu et al., 2014). The phylogenetic tree constructed according to the amino acid sequences of 33 members showed that the homological differences between members are relatively significant. ESD1 shares the closest homology with OsOFP7 and is also close to OsOFP27 and OsOFP8 (Fig. 1B). qRT-PCR showed that the expression levels of *ESD1* in roots, stems and leaves were very low, and there was a parabolic curve at stages 3-8 of young panicle development. The expression levels at stages 3-6 of panicle development gradually increased, peaked at stage 6, followed by a plunge at stage 7, and reached the lowest point at stage 8 (Fig. 1C). The roots, stems and leaves at the panicle differentiation stage as well as the young panicles at stages 3-8 were used for *in situ* hybridization. The results showed that the strongest hybridization signals of *ESD1* presented at stage 6 of panicle development, which mainly distributed in the ovules (Fig. 1, D-H). The results of *in situ* hybridization were consistent with those of qRT-PCR, indicating that ESD1 functions in ovules at the panicle development process in rice.

**Figure 1.**
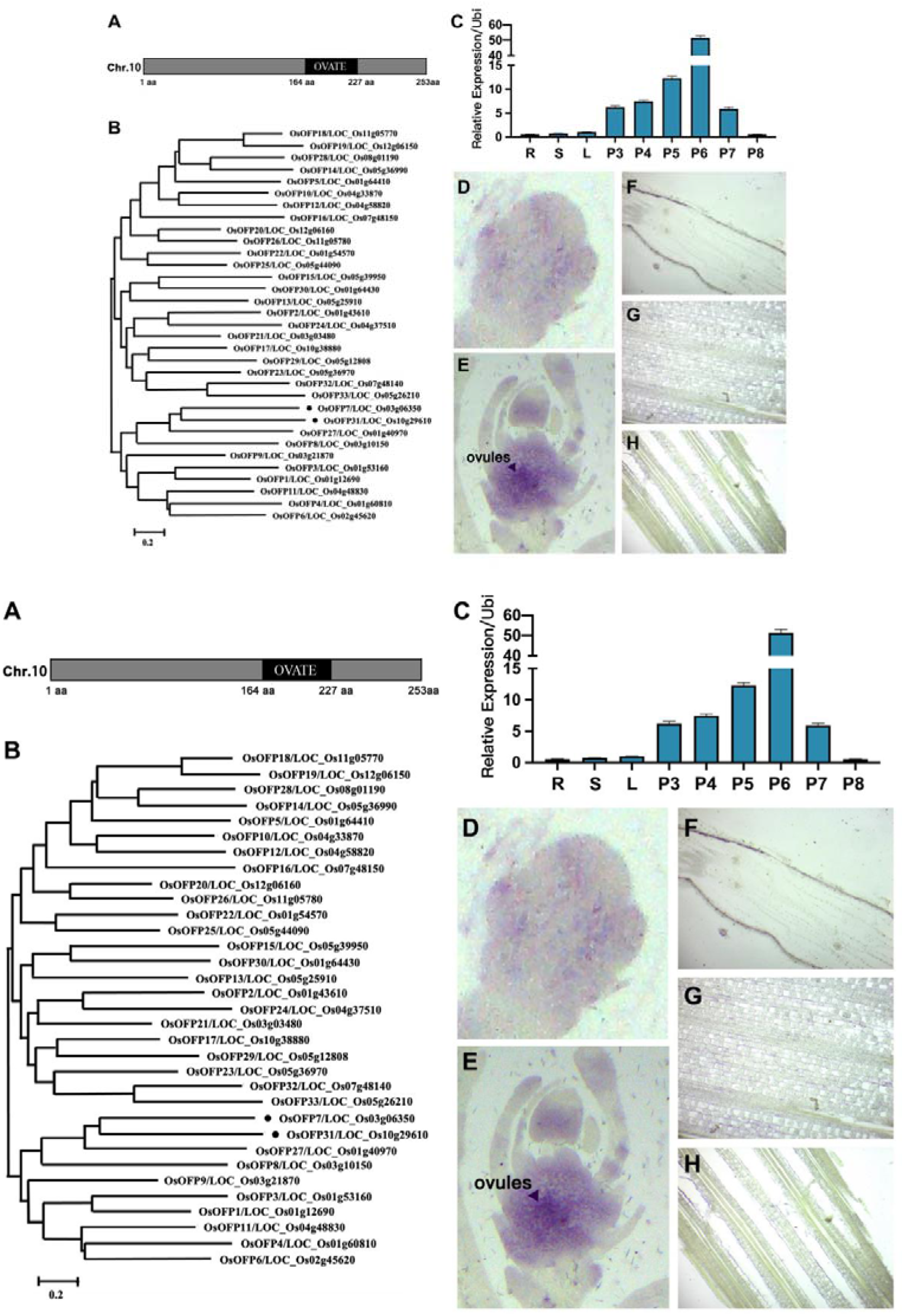
Gene structure and expression patterns of *ESD1.* A, Structural pattern diagram of *ESD1.* B, Phylogenetic tree of OFP family members. C, *ESD1* expressions in roots, stems, leaves, and young panicles at stages 3-8; R, roots; S, stems; L, leaves; P3-P8, young panicles at stages 3-8. D, *In situ* hybridization of *ESD1* at stage 3 of panicle development. E, *In situ* hybridization of *ESD1* at stage 6 of panicle development. F, *In situ* hybridization of *ESD1* in roots. G, *In situ* hybridization of *ESD1* in stems. H, *In situ* hybridization of *ESD1* in leaves.

### ESD1 predominantly localizes in cytoplasm

The recombinant vector *pGBKT7-ESD1* was transferred into yeast *(Saccharomyces cerevisiae)* strain Y2HGold. The strain expressing ESD1 proteins could not grow on the systematic tri-deficient medium (Fig. 2A), indicating that ESD1 was inactive in transcriptional activations. The *ESD1* and *GFP* fusion expression vector was transferred into rice protoplasts. The fluorescent signals of ESD1-GFP fusion proteins were mainly distributed in the cytoplasm (Fig. 2B).

**Figure 2.**
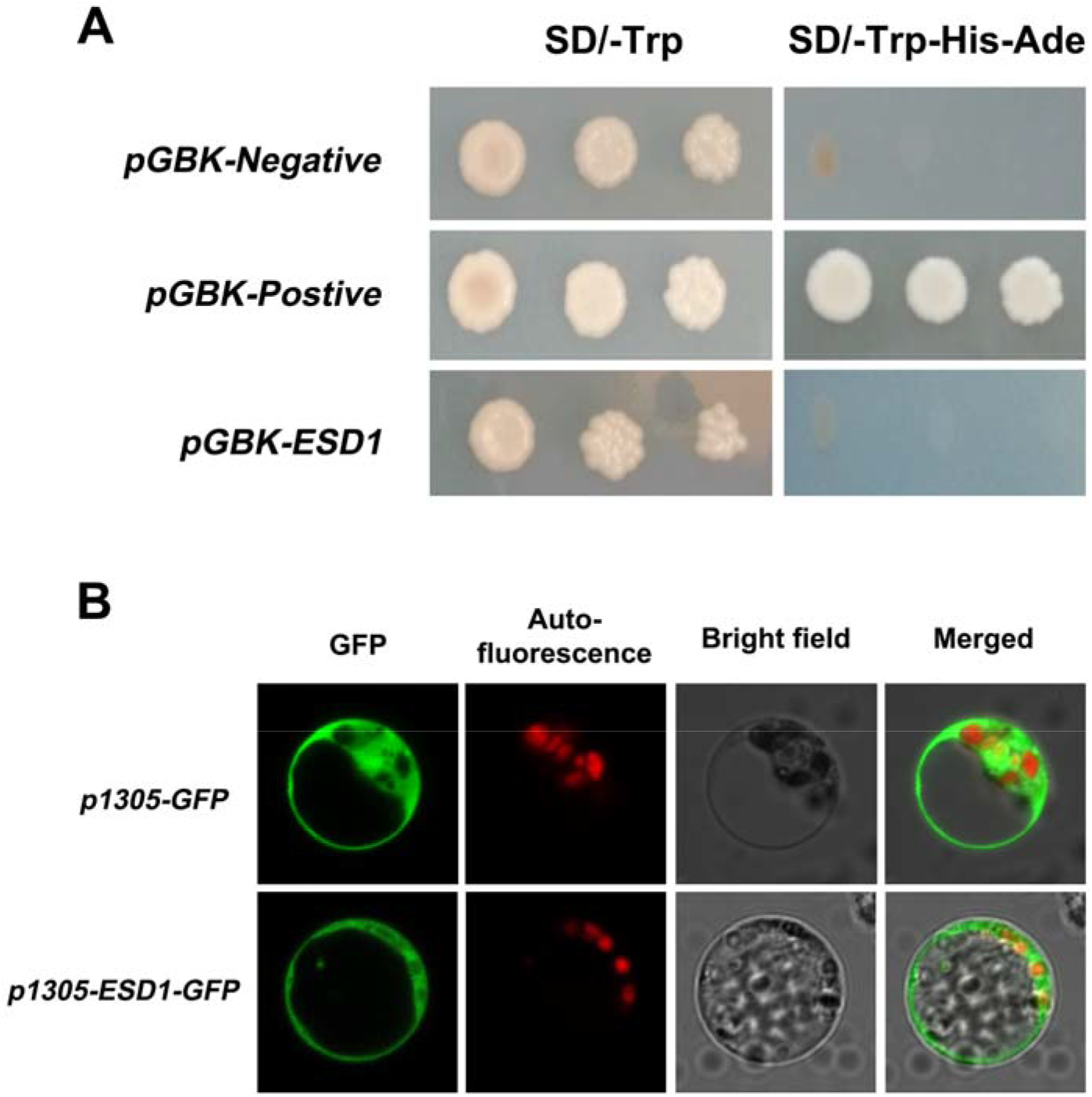
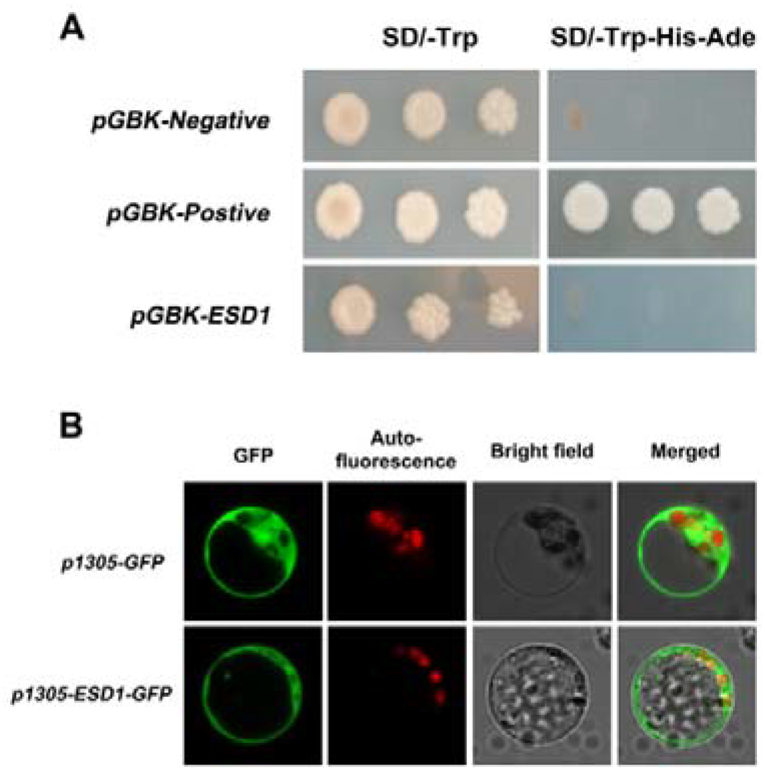
Protein subcellular localization of ESD1. A, Verification of ESD1 transcription activation. B, Subcellular localization of ESD1 in rice protoplasts.

### *esd1* mutants displayed a phenotype with reduced seed setting rates

In this study, two targeted editing sites (TS1 and TS2) of *ESD1* were constructed onto the CRISPR/Cas9 based binary expression vector (Fig. 3A). The calli induced from a *japonica* rice 9522 were infected with the *Agrobacterium-mediated* transformation process and subjected to tissue culturing, resulting in two edited homozygous mutants *esd1-m1* and *esd1-m2* as well as a heterozygous mutant *esd1-m3* (Fig. 3, B and C). The mutant *esd1-m1* was inserted with G and T at TS1 and TS2, respectively, resulting in an alteration of translated protein sequence after 71 aa, and an early termination after 91 aa, which caused the missing structure of the OVATE domain. The mutant *esd1-m2* was inserted with A at TS1 and deleted a sequence of AGTCGATGA at TS2, resulting in a change of translated protein sequence after 72 aa and an early termination after 91 aa, which also caused the missing structure of the OVATE domain. As for the heterozygous mutant *esd1-m3, ESD1* of one chromosome was normal, while *ESD1* of the other homologous chromosome was inserted with G at TS2, resulting in the translation of two types of protein, one of which was normal ESD1 protein, the other changed after 174 aa, which also caused the missing structure of the OVATE domain (Fig. 3D).

**Figure 3.**
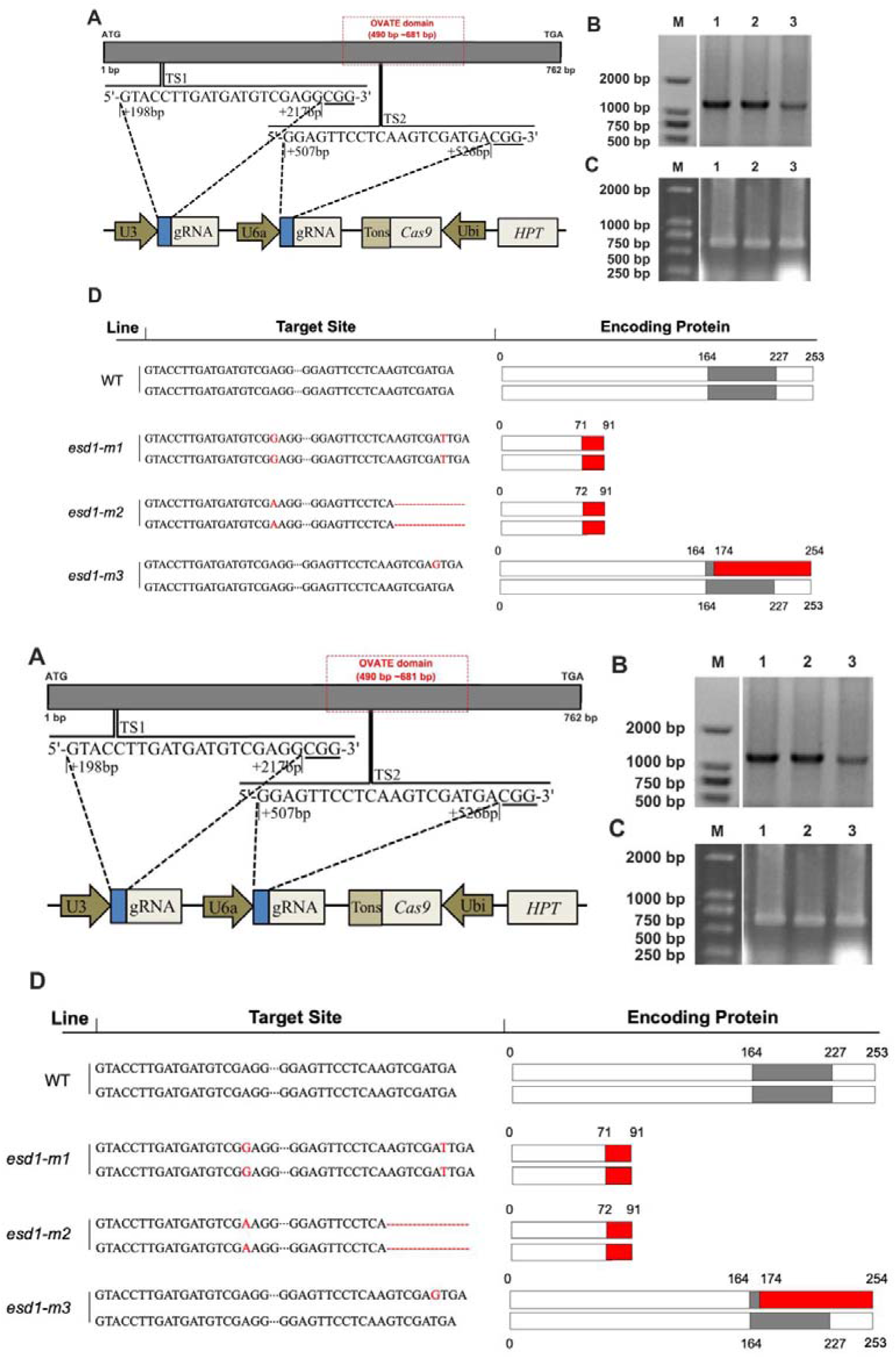
Targeted editing sites and types of mutations. A, Two targeted editing sites, TS1 and TS2, and information on CRISPR/Cas9 recombinant vector element. B, Screening of transgenic plants that integrate gene edited related sequence of the recombinant vector into the 9522 genome by PCR. C, Amplify the TS1 and TS2 sequences of transgenic plants by PCR. D, *ESD1’s* mutation types and translated protein residues. The grey box represents the OVATE domain, the red box represents the new amino acid sequence.

Phenotypic analysis indicated that no obvious difference was observed in the plant morphologies and internal spikelet structure between the wild type 9522 and the mutants *esd1-m1* and *esd1-m2* (Fig. 4A). Similar to 9552, I_2_-KI staining assay showed that the anthers of *esd1-m1* and *esd1-m2* were fertile (Fig. 4, C-H). Plant height, spikelets per panicle, 1000-grain weight, and tiller number of *esd1-m1* and *esd1-m2* were not significantly different from those of 9522 (Fig. 4, I, J, L, and M). However, *esd1-m1* and *esd1-m2* showed a phenotype with decreased seed setting rates, 53% and 58%, respectively, while that of the 9522 was 84% (Fig. 4, B, K), indicating that the deficiency of *ESD1* mainly reduced the spikelet fertility in rice. Meanwhile, the grain yield per plant of *esd1-m1* and *esd1-m2* also markedly decreased compared to that of 9522 (Fig. 4N). Whereas, no significant difference was detected in seed setting rates between *esd1-m3* and 9522. The population of T_1_ generation of *esd1-m3* showed a separation with seed setting rate. The result of co-segregation experiment showed that 86 single plants (heterozygous mutants and wild type plants) had normal seed setting rates, while 28 single plants (homozygous mutants named *esd1-m3h)* had reduced seed setting rates (Supplemental Fig. S1). Therefore, the phenotype with seed setting rate was controlled by *ESD1*.

**Figure 4.**
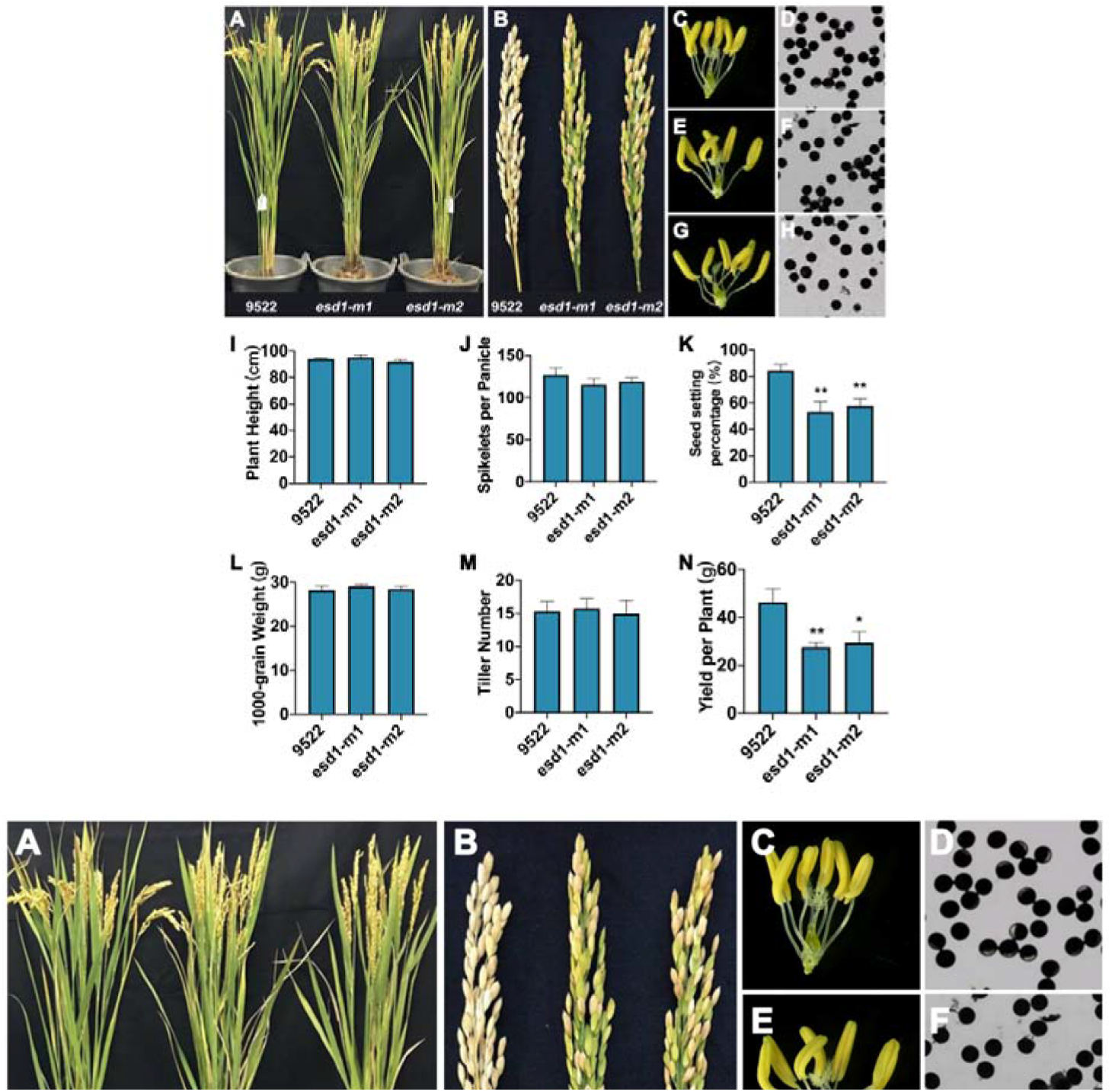
Phenotype identification of *esd1* mutants. A, Plant morphology of *esd1* mutants. B, Panicle types of *esd1* mutants. C, Anthers of 9522. D, Microscopy of 9522 pollen grains. E, Anthers of *esd1-m1.* F, Microscopy of *esd1-m1* pollen grains. G, Anthers of *esd1-m2.* H, Microscopy of *esd1-m2* pollen grains. I, Plant height of *esd1* mutants. J, Spikelets per panicle of *esd1* mutants. K, Seed setting rate of *esd1* mutants. L, 1000-grain weight of *esd1* mutants. M, Tiller number of *esd1* mutants. N, Yield per plant of *esd1* mutants. Asterisks indicate significant difference by Student’s *t* test (\**P*<0.05 and \*\**P*<0.01).

### Stamen development and pollen tube growth of *esd1* mutants were normal

Spikelet samples were taken at stage 8 of panicle development from 9522, *esd1-m1* and *esd1-m2*, and fixed with FAA fixative to prepare paraffin sections. The results showed that 9522, *esd1-m1* and *esd1-m2* had no significant differences in the structure of anthers, and all pollen grains were fertile (Fig. 5, A-C). Pollen grains were collected from 9522, *esd1-m1* and *esd1-m2* before flowering and cultured *in vitro.* The results showed that the germination rate of 9522 was 73.1%, while those of *esd1-m1* and *esd1-m2* were 72.0% and 71.1%, respectively, indicating that no significant differences was detected in the germination rates among the three genetic lines (Fig. 5, D-F). Spikelet samples at different stages after flowering were collected from 9522, *esd1-m1* and *esd1-m2.* The elongation process of pollen tubes was observed using an aniline blue staining. The results showed that the anthers of 9522, *esd1-m1* and *esd1-m2* could split open normally upon maturity. Their pollen grains could normally adhere to the stigma (Fig. 5, G-I), germinate pollen tubes along the styles, and eventually enter the embryo sac through the ovule hole (Fig. 5, J-L). These results indicated that the stamen development and pollen tube growth of *esd1* mutants were normal.

**Figure 5.**
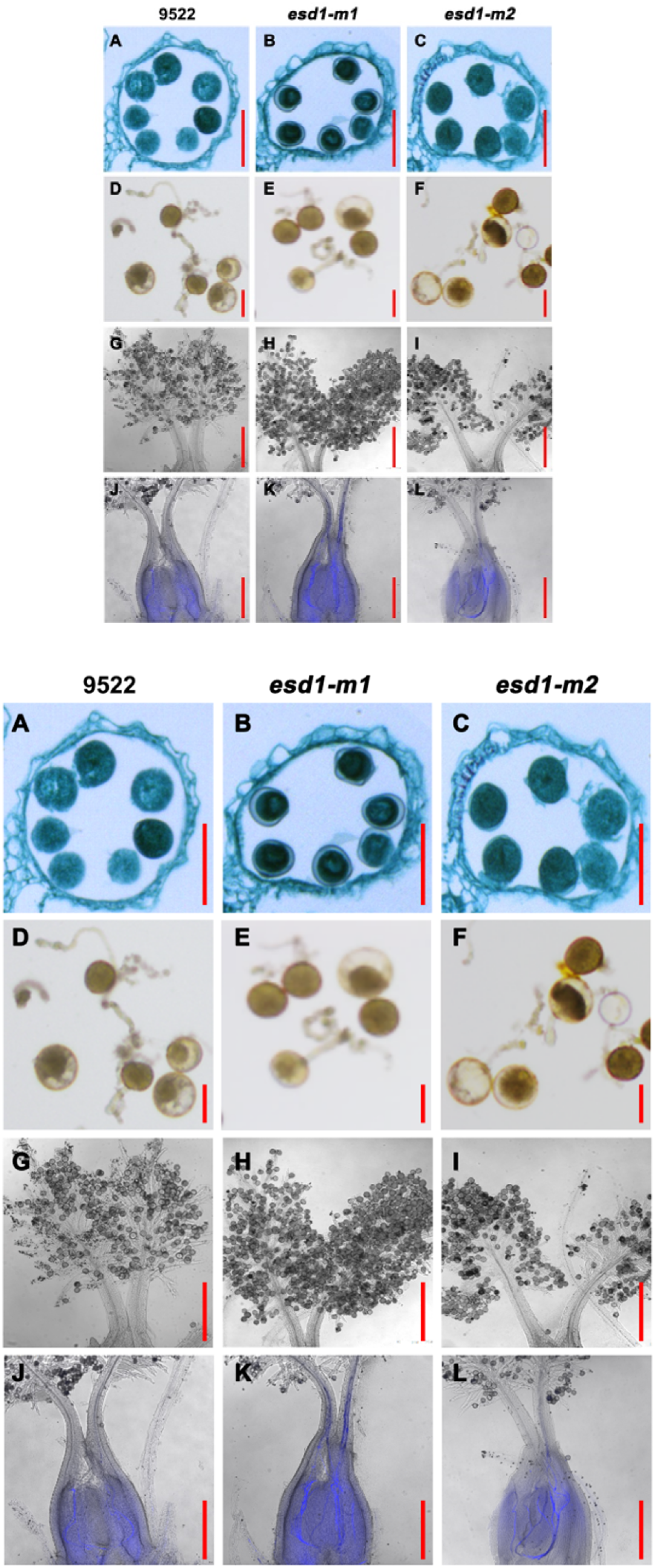
Microscopy of pollen grains and pollen tubes of *esd1* mutants. A to C, Anther paraffin sections of 9522, *esd1-m1* and *esd1-m2.* D to F, Pollen grain germinations *in vitro* of 9522, *esd1-m1* and *esd1-m2.* G to I, Pollen grains of 9522, *esd1-m1* and *esd1-m2* attached to the stigmas. J to L, Pollen tube growth of 9522, *esd1-m1* and *esd1-m2* (bright field microscopy + fluorescence microscopy).

### The pistil group of *esd1* mutants was abnormal

I_2_-KI staining was performed on the spikelets of 9522, *esd1-m1* and *esd1-m2* after flowering, and transparent xylene was used to examine the fertilization of spikelets. The results showed that the fertilization rate of 9522 was significantly higher than those of *esd1-m1* and *esd1-m2* (Fig. 6, A-D). The main panicles of *esd1-m1* were then subjected to natural pollination and saturated pollination. The results showed that the seed setting rate of *esd1-m1* natural pollination and saturated pollination were 61% and 64%, respectively, which have no significant difference (Fig. 6E). When the panicles of de-tasseled 9522’s main panicles were pollinated with *esd1-m1* pollens, the resulted seed setting rate of 9522 was 81%. However, when panicles of de-tasseled *esd1-m1’s* main panicles were pollinated with 9522 pollens, the seed setting rate of *esd1-m1* was 54% (Fig. 6F). These results further indicated that the anther development of *esd1* mutants was normal, and the cause of decreased seed setting rate of *esd1* mutants was affected by pistil group and occurred before fertilization.

**Figure 6.**
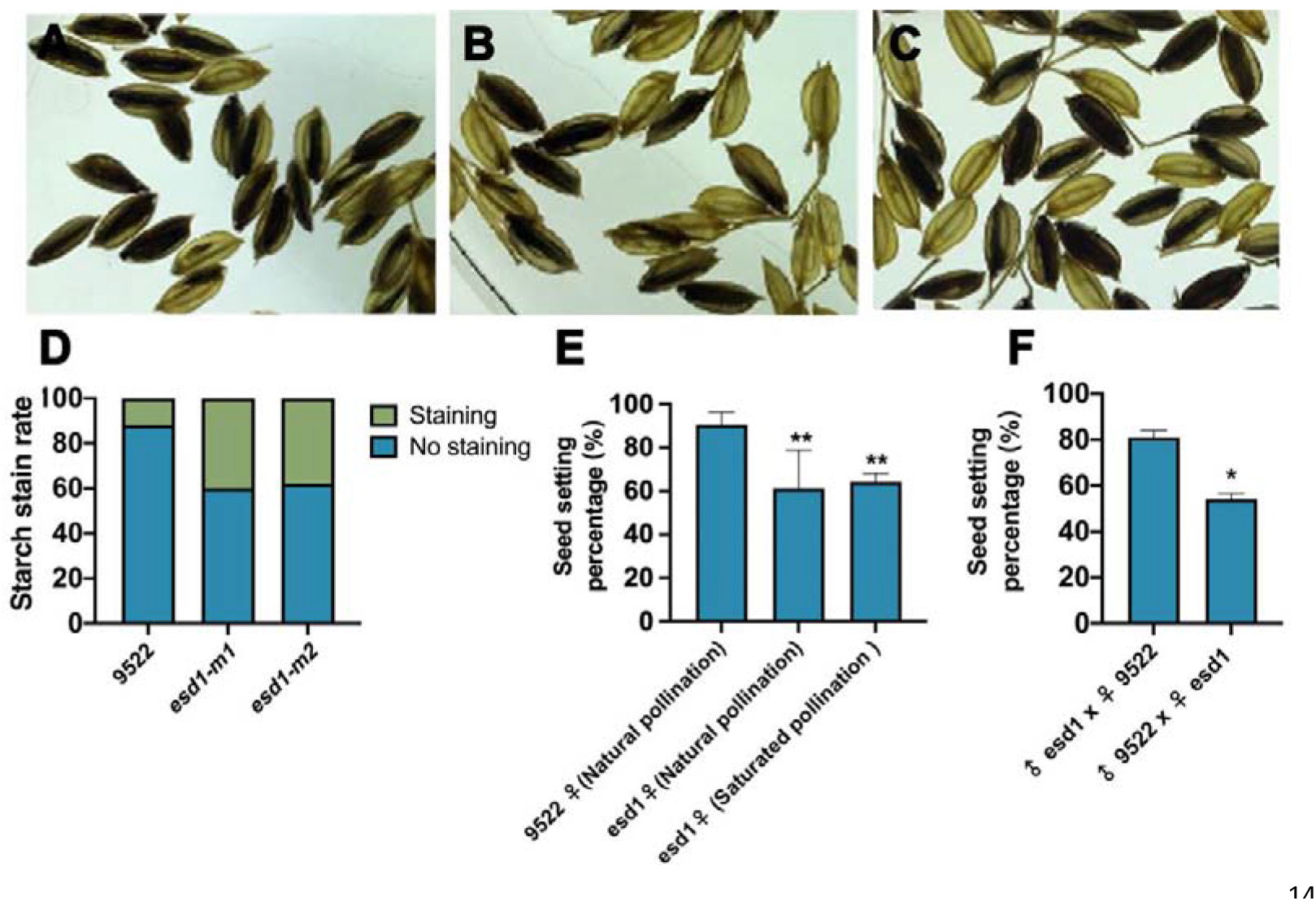
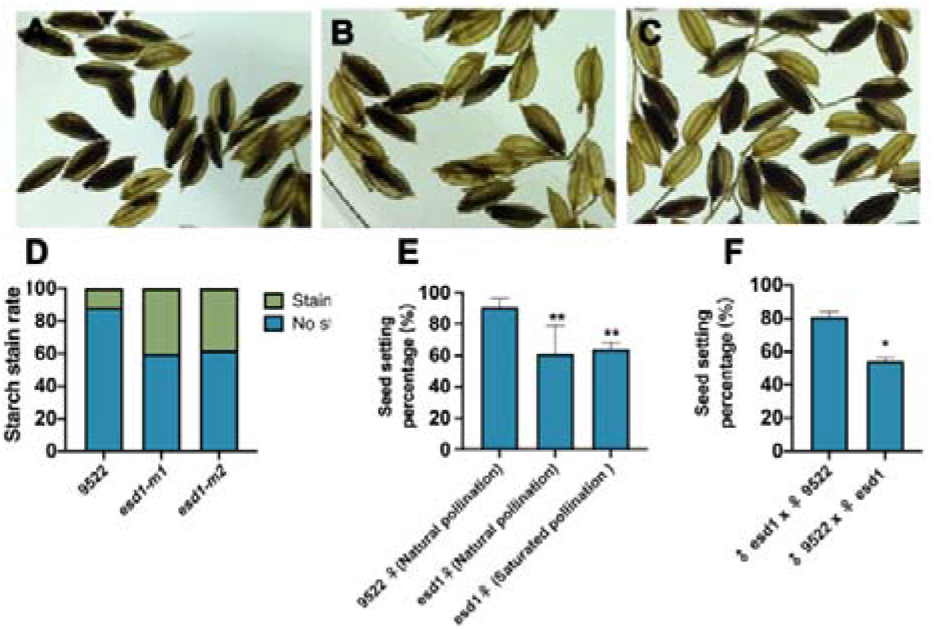
Fertilization and hybridization detection of *esd1* mutants. A to C, Embryo sac fertilization results of 9522, *esd1-m1* and *esd1-m2.* D, Starch stain rate of 9522, *esd1-m1* and *esd1-m2.* E, Natural pollination of *esd1* mutants and saturation pollination of *esd1* mutants with 9522 pollens. F, Reciprocal crosses between 9522 and *esd1-m1.* Asterisks indicate significant difference by Student’s *t* test (\**P*<0.05 and \*\**P*<0.01).

### No egg cells were detected in the partially matured embryo sacs of *esd1* mutants

A large number of mature embryo sacs of 9522, *esd1-m1* and *esd1-m2* were stained with nuclear-specific fluorescent dyes and hyalinized using hyalinization techniques. Samples were then scanned and photographed at 2 μm/layer under a confocal laser scanning microscope. The mature embryo sacs of 9522 were tested first, and 92% of their embryo sacs were found to develop healthily with a “seven cells and eight nuclei” structure at different levels of definition (Fig. 7, A and D). The mature embryo sacs of *esd1-m1* were then observed, and 58% of their embryo sacs developed normally, whereas 42% failed to show any egg cells and synergids despite the existence of normal antipodal cells and central polar nuclei (Fig. 7, B and D). Similarly, among the embryo sacs of *esd1-m2,* 56% developed normally, whereas 44% showed no egg cells and synergids (Fig. 7, C and D). The transcriptional levels of genes involved in the development of rice embryo sac were detected in the young panicles of 9522 and *esd1* mutants by qRT-PCR. The results showed that the deficiency of *ESD1* could significantly reduce the expression of *OsAPC6* (Fig. 7E). These results demonstrated that abnormal development of some embryo sacs led to a reduction in seed setting rates in *esd1* mutants.

**Figure 7.**
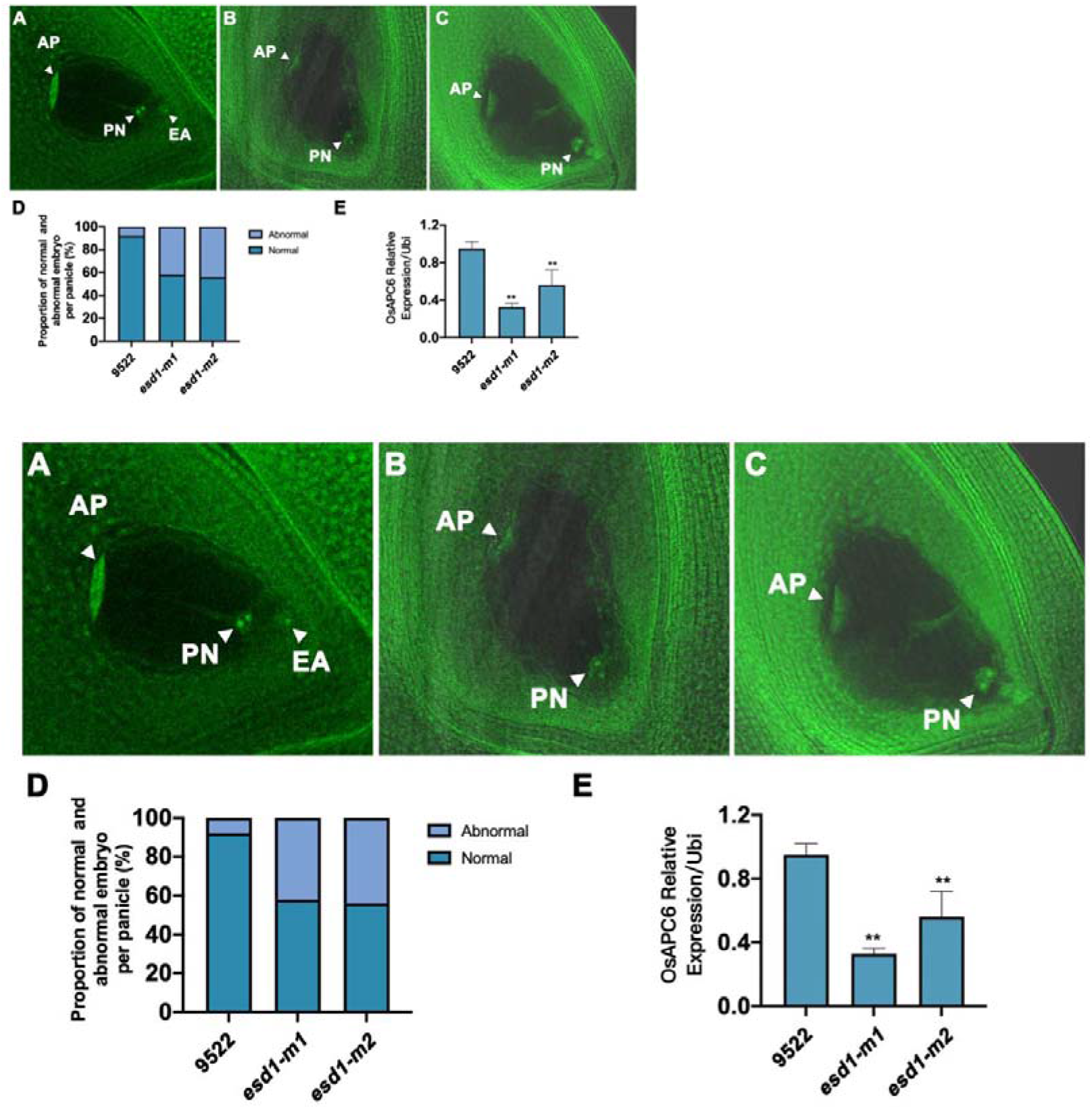
Detection of mature embryo sac of *esd1* mutants and related qRT-PCR. A, Normal embryo sac of 9522; AP, antipodal cells; PN, polar nuclei; EA, egg cell. B, Abnormal embryo sac of *esd1-m1.* C, Abnormal embryo sac of *esd1-m2.* D, Abnormal embryo sac ratio of *esd1* mutants. E, Expression of *OsAPC6* in the young panicles of 9522 and *esd1* mutants. Asterisks indicate significant difference by Student’s *t* test (\*\**P*<0.01).

### The ubiquitin-mediated protein degradation pathway was the most involved metabolic pathway in *esd1* mutant differentially expressed genes

The young panicles at stages 5 and 6 of panicle development of 9522 and *esd1-m1* were taken to transcriptome analysis. A total of 190 significant differentially expressed genes (DEGs) were detected, and 51 common DEGs were presented at two stages. One hundred and two significant DEGs were detected in *esd1-m1* with 81 up-regulated and 21 down-regulated expression genes at stage 5. A total of 88 significant DEGs, 72 up-regulated and 16 down-regulated genes, were detected in *esd1-m1* at stage 6 (Fig. 8A). The results of hierarchical cluster analysis showed that DEGs of 9522 and *esd1-m1* could cluster together at stage 5, indicating a high degree of similarity in genetic background and gene expression patterns between 9522 and *esd1-m1* (Fig. 8B). One hundred and ninety DEGs were classified into eight functional categories, namely ubiquitin mediated proteolysis, circadian rhythm, biosynthesis of amino acids, nucleotide excision repair, flavonoid biosynthesis, cysteine and methionine metabolism, RNA transport and ribosome. The ubiquitin-mediated protein degradation pathway is the most DEGs involved metabolic pathway (Fig. 8C). Meanwhile, the transcriptome analysis also showed that the transcriptional level of *OsAPC6* was significantly reduced in *esd1-m1*, which was highly consistent with the result of qRT-PCR (Fig. 7E). *OsAPC6* was identified to encode a cell cycle-related ubiquitin ligase that degrades mitosis-related regulators via the ubiquitin-proteasome pathway (Kumar et al., 2010). These results imply that the ubiquitin-mediated protein degradation pathway might be involved in regulating the development of embryo sacs in rice.

**Figure 8.**
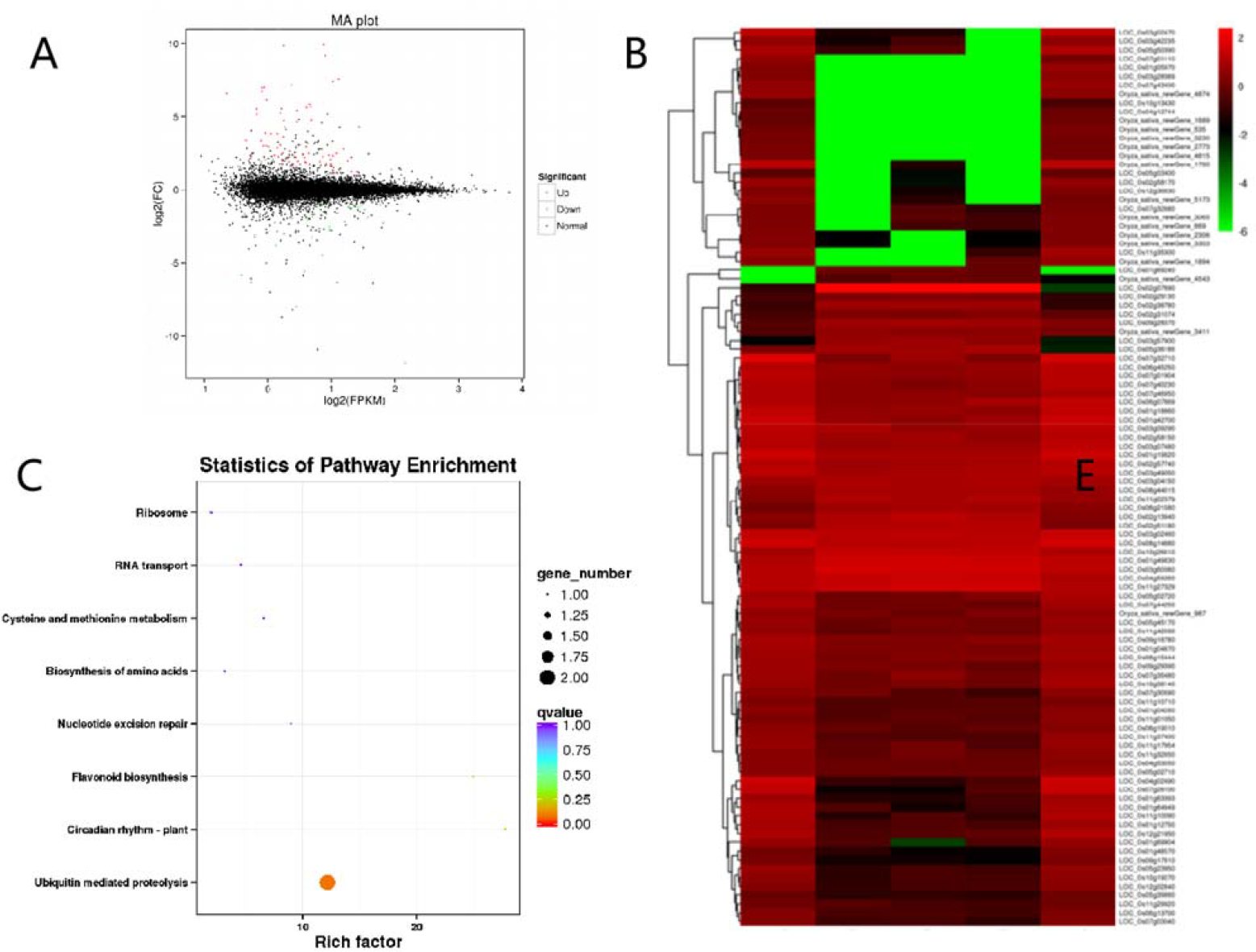
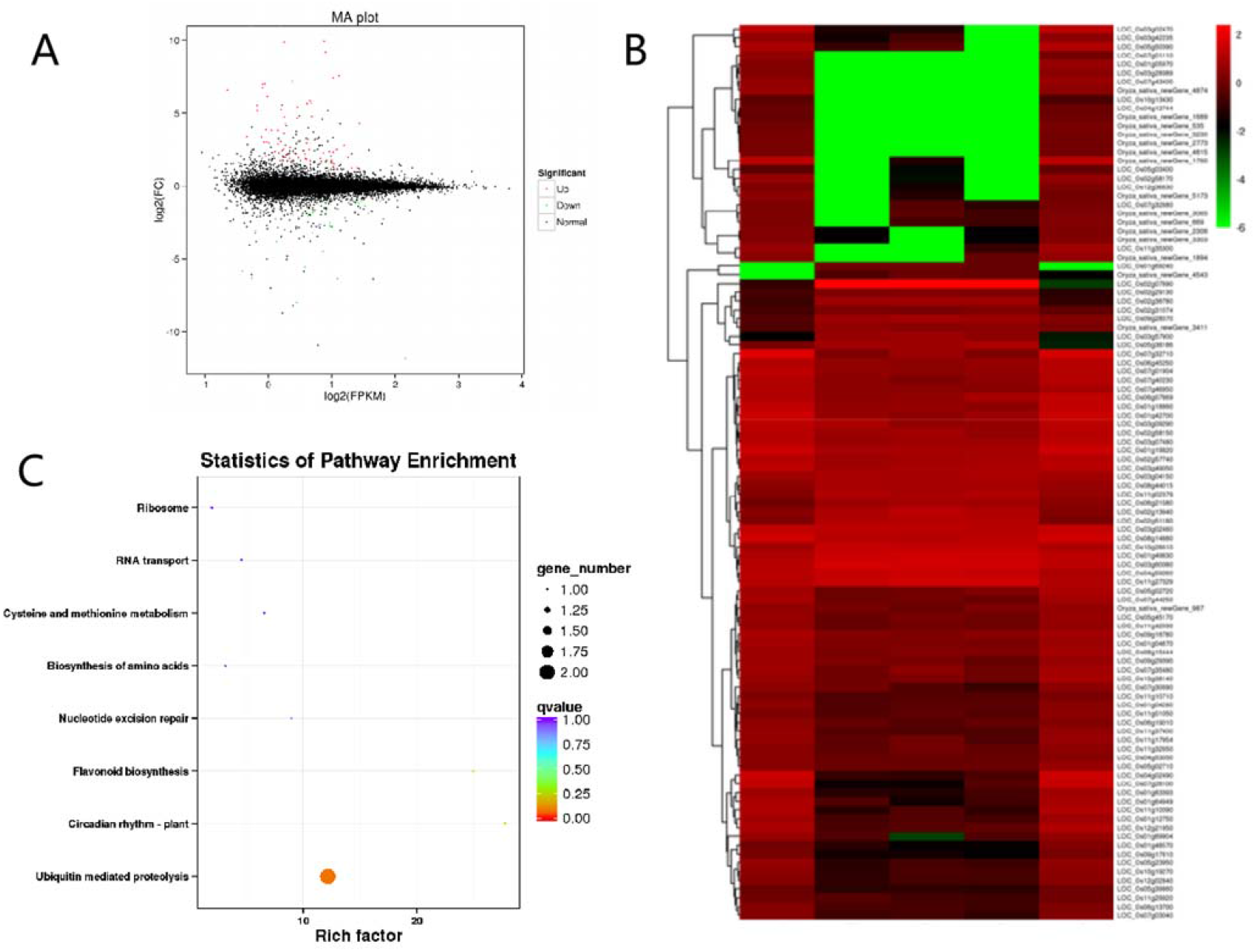
Transcriptome analysis of *ESD1.* A, Differential expression volcano map. B, Differentially expressed gene cluster map. C, Scatter plot of enrichment of differentially expressed gene KEGG pathway.

## Discussion

Seed setting rate, grain number and grain weight are the most important factors that determine rice yield. Seed setting rate is affected by several factors including the pistil and stamen development. It has been reported that OFP family members can regulate pistil development. For example, the OFP family member AtOFP5 protein is involved in regulating ovule formation in Arabidopsis (Hackbusch et al., 2005; Pagnussat et al., 2007). In our previous study, microarray data showed that ESD1 protein which contains an OVATE domain located at 164-227 aa and belongs to the OFP family was highly and abundantly expressed mainly in spikes. In this study, we obtained several different types of loss-of-function mutants of *ESD1* by CRISPR/Cas9 technology (Ma et al., 2016; Sun et al., 2016; Zhang et al., 2018; Selma et al., 2019; Zhang et al., 2019; Ma et al., 2020). The seed setting rates of *esd1* mutants were significantly lower compared to that of wild type 9522, and the co-segregation test showed that the trait was controlled by a single gene. The above research indicated that *ESD1* is a key modulator of rice seed setting rate.

Low seed setting rate in rice can be caused by many factors (Zhou et al., 2011; Li et al., 2013; Xu et al., 2019). The pollen fertility I_2_-KI staining, anther paraffin sections, *in vitro* culture of pollen grains and pollen tube germination test *in vivo* all proved that the stamen development and pollen tube growth of *esd1* mutants were normal. These results suggest that the reduced seed setting rate of *esd1* mutants is not resulted from the development of stamens. Promoter element prediction showed that *ESD1* has elements involved in photoresponse, such as ACE, Box 4, G-box, G-Box and GA-motif, *etc*. (Li et al., 2017). To demonstrate whether the fertility of the *esd1* mutants were affected by the environment, the *esd1* mutants were planted under different photoperiods and temperature treatments. The results showed that anther developments of 9522 and *esd1* mutants were normal under different photoperiods and temperature treatments (Supplemental Fig. S2, S3), and no significant difference was detected in the seed setting rates of *esd1* mutants under various treatments, indicating that the functions of *ESD1* were not affected by light or temperature.

The results of embryo sac fertilization, reciprocal cross test, natural pollination and saturation pollination further demonstrated that the anther development of *esd1* mutants was normal, and the reduced seed setting rates of *esd1* mutants was resulted from the development of pistil group before fertilization. Mature embryo sac structure inspection proved that the embryo sac abortion of *esd1* mutants was caused by the absence of egg cell. The presence of normal polar nuclei and antipodal cells in all aborted embryo sacs further suggested that embryo sac development was normal during the meiosis from megasporocyte and first two mitotic divisions from functional macrospore. During or after the third mitosis of functional megaspores, egg cells and synergids gradually degenerate, and this process is not influenced by light and temperature. The results of both qRT-PCR and *in situ* hybridization demonstrated that *ESD1* expression was highest at the 6^th^ stage of panicle differentiation. Stage 6 is the time when functional megaspores begin to undergo mitosis (Liu et al., 1997). The alteration of *ESD1* expression level coincided with the failure of embryo sacs of *esd1* mutants, which further indicated that *ESD1* regulates embryo sac development by influencing mitosis of functional megaspores.

Transcriptome data analysis revealed differentially expressed genes in *esd1* mutant and wild type material. The ubiquitin-mediated protein degradation pathway was the most involved metabolic pathway in differentially expressed genes in *esd1* mutants. The ubiquitin-proteasome system is the major pathway of intracellular protein degradation, involved in more than 80% of protein degradation (Tokumaru et al., 2017; Zhang et al., 2017; Xia et al., 2020; Majumdar et al., 2020). The intracellular or extracellular ubiquitin proteasome system is likely to be involved in regulating the development of embryo sac by acting on different substrates (Liu et al., 2008; Kumar et al., 2010; Awasthi et al., 2012). Transcriptome analysis showed that the deficiency of *ESD1* resulted in the down-regulation of *OsAPC6* encoding a cell cycle-related ubiquitin ligase involved in the degradation of mitosis-related regulators via the ubiquitin-proteasome pathway, which was consistent with the qRT-PCR result. In the T-DNA insertion mutant of *osapc6,* the functional megaspores in some of embryo sacs showed an abnormality during the second mitosis, therefore the seed setting rate of *osapc6* mutants was reduced by 45%. Coincidentally, the seed setting rate of the *esd1* mutants also decreased by about 45%. It is hypothesized that ESD1 has a genetic regulatory relationship with OsACP6, both act simultaneously during mitosis of functional macrospores. Consequently, ESD1 might have some regulatory interactions with OsAPC6, which could degrade relevant enzymes involved in the mitosis of functional megaspores through the ubiquitin-proteasome pathway, promote the programmed death of egg cells, and eventually lead to reduction in seed setting rate. However, more data and in-depth studies are needed on how ESD1 affects the ubiquitination pathway and thus regulates the development of the embryo sac.

Subcellular localization analysis revealed that ESD1-GFP fusion proteins predominantly localized in the cytoplasm (Fig. 2B). There are 33 genes that encode the full-length OVATE domain containing proteins in the rice genome. Some of the OFP family members are generally located in the nuclei and function as transcription factors (Wang et al., 2008; Schmitz et al., 2015). Ovate family proteins are not a typical family of transcription factors. For example, the Arabidopsis ovate family proteins (AtOFPs) have been shown to function as transcriptional repressors (Wang et al., 2016c). In this study, we performed subcellular localization prediction for ESD1 protein with TargetP tools (http://www.cbs.dtu.dk/services/TargetP/), and the result showed that ESD1 does not contain nuclear localization signal sequences (Supplemental Table S1). In addition to the subcellular localization study in rice protoplasts, we also repeated this experiment in tobacco, and the results showed that the fluorescent signal was distributed throughout the cell, including the nucleus and cytoplasm (Supplementary Figure S4), which indicated that ESD1 does not contain the typical nuclear localization signal peptide. Verification of ESD1 transcription activation showed that ESD1 was inactive in transcriptional activations. Therefore, we speculate that ESD1 might be an atypical transcription factor to regulate the development of embryo sac in rice, but a further investigation should be performed in the future study.

In conclusion, this study demonstrated that ESD1 positively regulates the seed setting rate by controlling embryo sac development in rice, and has implications for the improvement of rice yield. These results not only help to further illustrate the regulatory mechanism of embryo sac development but also have high theoretical value in revealing the molecular mechanism of rice grain formation.

## MATERIALS AND METHODS

### Phylogenetic analysis of ESD1

Protein sequence comparisons were conducted using MUSCLE v3.6 based on the BLAST given in TIGR (Rice Genome Annotation Project, http://rice.plantbiology.msu.edu/index.shtml) and PlantTFDB (Plant Transcription Factor Database, http://planttfdb.cbi.pku.edu.cn) of OFP family members. A phylogenetic tree was constructed with the aligned relative protein sequences using MEGA v3.0 (http://www.megasoftware.net/index.html) and Neighbor-Joining method with the following parameters: poisson correction, pairwise deletion, and bootstrap (1,000 replicates; random seed).

### Construction and transformation of CRISPR/Cas9 vectors

We used CRISPR/Cas9 target site designing website called E-CRISP (http://www.e-crisp.org/E-CRISP/) to analyze and design the *ESD1* sequences. Target sites TS1 and TS2 with high scores were selected to reduce the probability of missing shots. TS1, TS2, and their corresponding reverse complementary sequences were then made into 2 pairs of primers, *TS1-U3-F/R* and *TS2-U6a-F/R* (ggca and aaac were bound at the 5’ ends of the forward primer and reverse primer respectively; Supplemental Table S2 for primer sequences). Through Golden Gate cloning, *DS-TS1* and *DS-TS2,* the double-stranded sequences obtained by annealing the above two pairs of primers, were inserted into the intermediate vectors *pYLgRNA-U3* and *pYLgRNA-U6a,* respectively. Two expression cassettes *U3-TS1-gRNA* and *U6a-TS2-gRNA* were obtained via PCR amplification (Supplemental Table S3 for the amplification primer sequences). Through Golden Gate cloning, two expression cassettes were then inserted into the plant expression vector *pYLCRISPR/Cas9-MH,* after seedlings were obtained by tissue culture, primers SP1/SP2 were used to detect whether *pYLCRISPR/Cas9-MH* was introduced into the recipient plants (Supplemental Table S4 for primer sequences), and then primers *ESD1* -JC-F/R were used to amplify sequences containing the target sites TS1, TS2 (Supplemental Table S4 for the amplification primer sequences), and then sequencing was used to detect the mutations at the target sites.

### Microscopy of pollen grains

Spikelet samples of 9522, *esd1-m1,* and *esd1-m2* were collected from their main panicles (in the same position) 1 day before flowering, fixed in Kano solution (absolute ethanol: glacial acetic acid=3: 1), and stored at 4°C. For microscopic examination, each spikelet was stained with 1% I2-KI and observed under a microscope with a 10 X ocular. The field of view was switched at least 3 times. The total count of pollen grains was greater than 500. Judging criteria: the round dark brown pollen grains were considered fertile; irregularly shaped, weakly stained or unstained pollen grains were deemed sterile.

### Anther paraffin sections

Spikelet samples at stage 8 of panicle development were taken from 9522, *esd1-m1*, and *esd1-m2*. Samples were dehydrated, hyalinized, immersed, and embedded in paraffin. Samples were then cut into 10 μm-thick sections, which were stained with safranin O-fast green, hyalinized, and sealed. The sections were observed under an Olympus BH2 optical microscope.

### Pollen grain culture *in vitro*

An appropriate amount of liquid culture medium was placed on glass slides. When the plants were about to bloom, the spikelets of 9522, *esd1-m1,* and *esd1-m2* were pinched with tweezers, after which their pollen grains were gently shaken off onto liquid media, respectively. The glass slides were covered with coverslips and placed in petri dishes with moistened filter paper. The petri dishes were sealed and incubated at 28°C in the dark for 60 min. Afterward, a drop of 0.005% water-soluble aniline blue solution was added to each liquid culture medium, and the germination of pollens was checked under a fluorescence microscope. The above procedures were repeated three times for each group of samples. More than 200 pollen grains were detected each time. The average percentage of germinated pollens was deemed as the germination rate.

### Pollen tube germination test *in vivo*

After heading, the flowering time of 9522, *esd1-m1,* and *esd1-m2* was written down with a marker. Spikelet samples were collected at 0, 10, and 20 min, and 1, 1.5, 2, and 2.5 h after flowering, fixed with FAA fixative for 24 h, immersed in 70% ethanol solution and stored at 4°C. The ovaries were peeled off before observation. The samples were then rehydrated in 50%, 30%, and 10% ethanol (30 min each), washed with ddH_2_O two to three times, immersed in 2 mol/L NaOH solutions to soften for 12 h, rinsed with ddH_2_O several times, stained with 0.05% aniline blue for 12 h, and then observed under a confocal laser scanning microscope (Zeiss LSM 880).

### Saturation pollination

One day before flowering, five individual plants sharing the same growth status among the *esd1-m1* population were bagged. There were 2 panicles per bag and 2 bags per plant, one of which was used for saturation pollination, and the other for self-pollination. For saturation pollination, *esd1-m1* plants were pollinated with 9522 pollens at the full-bloom stage. This process was repeated for seven consecutive days. Seeds were tested upon harvest, and the seed setting rate of saturation pollination and self-pollination were compared.

### Reciprocal cross tests

The main panicle samples of 9522 and *esd1-m1* were collected at the heading stage. Bloomed spikelets and spikelets that could not bloom on that day were cut off. The spikelets that could bloom on that day were retained and de-tasseled before flowering. At the full-bloom stage, de-tasseled spikelets of the *esd1-m1* main panicles were pollinated with 9522 pollens, and de-tasseled spikelets of the 9522 main panicles were pollinated with *esd1-m1* pollens. Seeds were tested upon harvest, and the seed setting rate of the two hybrids were compared (Li et al., 2011b).

### Tests for fertilization

Spikelet samples were taken from 9522, *esd1-m1,* and *esd1-m2* one day after flowering, and immersed in 40°C water overnight, followed by 35% concentrated hydrochloric acid for 30 min. The residual hydrochloric acid was rinsed off with ddH2O. The samples were then immersed in the I2-KI solution for an additional 30 min. After rinsing off the excess iodine with 70% alcohol, the samples were immersed in absolute ethanol for 30 min. Lastly, the spikelets were hyalinized in xylene and examined on a fluorescent box.

### Mature embryo sac structure inspection

Spikelet samples at stage 8 of panicle development were collected from 9522, *esd1-m1*, and *esd1-m2*, fixed with FAA fixative, and then immersed in 70% ethanol after ovaries were peeled off. In this process, the ovaries of different samples should be respectively immersed in ethanol at gradient concentrations (50%, 30%, and 15%) for 20 min each, and rinsed with ddH2O 2 to 3 times. After that, the ovaries were hydrolyzed with 1 mol/L HCl in a 60°C water bath for 15 min. Subsequently, the ovaries were stained with 1% eosin Y for 8 h. After staining, the ovaries were rinsed with ddH_2_O until colorless and pre-treated with 0.1 mol/L citric acid-disodium hydrogen phosphate buffer (pH=5.0) for 8 h. They were then dyed with 20 μg/ml H33342 in the dark at 25°C for 24 h. The ovaries were rinsed with ddH2O 2 to 3 times and dehydrated in ethanol at gradient concentrations (15%, 30%, 50%, 70%, 85%, and 95%) for 20 min each. The ovaries were then dehydrated twice with absolute ethanol for 2 h each time and transferred into absolute ethanol overnight. The next day, the dehydrated ovaries were immersed in a mixture of equal amounts of methyl salicylate and absolute ethanol for 1 h as a transition and then hyalinized with methyl salicylate three times. The first two times took 2 h each, and the last time took 15 h, after which the ovaries could be stored in methyl salicylate. A drop of sesame oil was placed on each concave glass slide before observation, and a treated ovary was then mounted on a slide. After that, the slide was inverted under a confocal laser scanning microscope and scanned at 2 μm/layer under 488 nm laser. Meanwhile, photographs were taken.

### qRT-PCR

Total RNA was extracted using the TRIzol method, and cDNA was obtained by reverse transcription after digesting the RNA. The qRT-PCR reaction system (ABI QuantStudio 3) included: *ESD1*-qP-F/R primers (Supplemental Table S5 for primer sequences), 0.5 μl each, 2 μl cDNA, 4.5 μl SYBR, 0.5 μl ROX, and 2.0 μl ddH_2_O. Three biological replicates and three technical replicates were taken. Settings for the qRT-PCR amplification program: 95°C for 5 min; 95°C for 15 s; 58°C for 35 s; 45 cycles; default settings for dissolution curve and cooling temperature. The internal reference standard was the expression level of *OsACTIN1,* and the expression of *ESD1* was calculated using the 2^-ΔΔCT^ formula (Livak et al., 2001).

### *In situ* hybridization

Roots, stems, leaves at the stage of panicle differentiation, and young panicles at stages 3-8 were taken (the whole process was guaranteed to be free of RNase contamination), fixed with FAA fixative, and then made into paraffin sections. *ESD1* -ish-F/R primers (Supplemental Table S6 for primer sequences) and *in situ* hybridization probes were designed for *in vitro* transcription. Kawata’s experimental method was used for reference in the sensitivity test of the *in situ* probes (Kawata et al., 2010); Kouchi’s experimental method was used for reference in the RNA hybridization and immunoassay (Kouchi et al., 1993).

### Subcellular localization in rice protoplast

*ESD1* and *GFP* fusion expression vector was constructed. A 10 μl of fusion expression vector plasmid and 110 μl of protoplasts were pipetted, placed in a 2 ml centrifuge tube, and mixed well. A 130 μl of 40% PEG4000 prepared on-site was added to the mixture, which was then let sit at room temperature for 15 min. 500 ml of W5 solution was added to stop the reaction, and the mixture was then centrifuged at 1188 rpm for 2 min, with the supernatant discarded. Then, 1 ml of W5 solution was added to re-suspend the protoplasts. The mixture was again centrifuged at 1188 rpm for 2 min, with the supernatant discarded. Finally, 1 ml of W5 solution (containing 1 μl of Kan) was added to re-suspend the protoplasts. The mixture was then transferred onto a 24-well plate, incubated at 28°C, 40 rpm in the dark for 12-16 h, and then observed under a confocal laser scanning microscope.

### Transcription activation verification

The *pGBKT7-ESD1* vector was constructed. *pGBKT7-CSA* vector plasmid (positive control) and *pGBKT7* empty vector plasmid (negative control) were used to transform Y2H Gold yeast competent cells using a heat shock approach. The bacterial solution was spread over the SD/-Trp solid medium, which was then sealed, inverted, and kept in an incubator at 28°C for 3-5 d. Once the colonies reached 2-3 mm, a single colony from each of the 3 solid media was picked and diluted at 1:10, 1:10^3^, and 1:10^5^. 4 μl of the diluents was transferred onto an SD/-Trp single-deficient plate and also onto an SD/-Trp-His-Ade triple-deficient plate. The plates were sealed, inverted, and kept in an incubator at 30°C for 3-5 d after air-drying the bacterial plaques.

### Transcriptome analysis

Extract and isolate mRNA from 9522, *esd1-m1* plant young panicles at 5 and 6 stages of development, add crushing buffer, interrupt the mRNA into short sequences (200-700 nt), and then reverse transcribe to obtain double-stranded cDNA; purified, eluted, with base A added to the end and connected to the sequencing junction; and then enrich the cDNA library by PCR, with Illumina HiSeq 2000 sequenced the library. The sequences obtained after sequencing and filtering were compared to the reference genome of the *Japonica* variety Nipponbare.

## Accession Numbers

Sequence data from this article for the cDNA and genomic DNA of *ESD1* can be found in the GenBank/EMBL/Gramene data libraries under accession number LOC_Os10g29610.

## Supplemental Data

The following supplemental materials are available.

**Supplementary Figure S1.** Phenotypic data of *esd1-m3h.*

**Supplementary Figure S2.** Effects of photoperiods on the fertility of *esd1* mutants.

**Supplementary Figure S3.** Effects of temperature treatments on the fertility of *esd1* mutants.

**Supplementary Figure S4.** The subcellular localization of ESD1 in tobacco.

**Supplemental Table S1.** Subcellular localization prediction for ESD1 protein with TargetP.

**Supplemental Table S2.** Amplification Primers for target sites TS1 and TS2.

**Supplemental Table S3.** Amplification primers for expression cassettes.

**Supplemental Table S4.** Detection primers for target sites mutation.

**Supplemental Table S5.** Primers for qRT-PCR reaction.

**Supplemental Table S6.** Primers for *in situ* hybridization.

## ACKNOWLEDGMENTS

We are grateful to Prof. Dabing Zhang, Shanghai Jiao Tong University for kindly providing the microarray data. We are grateful to Prof. Yaoguang Liu, South China Agricultural University for kindly providing the CRISPR/Cas9 editing vectors, *pYLgRNA-U3*, *pYLgRNA-U6a*, and *pYLCRISPR/Cas9-MH*.

